# Bioactivity-driven high throughput screening of microbiomes of medicinal plants for discovering new biological control agents

**DOI:** 10.1101/611855

**Authors:** Irum Iqrar, Zabta Khan Shinwari, Ashraf El-Sayed, Gul Shad Ali

## Abstract

In a preliminary DNA-based microbiome studies, diverse culturable and unculturable bacterial taxa were identified in the roots and rhizospheres of different medicinal plants. In this report, culturable endophytic bacteria were isolated from four economically important medicinal plants *Dodonaea viscosa, Fagonia indica, Caralluma tuberculata* and *Calendula arvensis.* On the basis of initial antimicrobial screening, nine bacterial species in seven different genera, *Streptomyces, Pseudomonas, Enterobacter, Bacillus, Pantoea, Pseudarthrobacter* and *Delftia*, were selected for further analyses. These bacteria were identified using 16S rRNA gene sequencing. Antimicrobial assays of these selected bacteria revealed that *Pseudomonas taiwanensis* has strong anti-*Phytophthora* activity. Volatiles produced by *P. taiwanensis* inhibited growth of *P. parasitica* more than 80%. Ethyl acetate extracts of *S. alboniger* MOSEL-RD3, *P. taiwanensis* MOSEL-RD23, *E. hormaechei* MOSEL-FLS1 and *B. tequilensis* MOSEL-FLS3 and *D. lacustris* MB322 also displayed high potency against *P. parasitica*. All these bacterial extracts showed strong inhibition against *P. parasitica* at different concentrations (4 µg/mL – 400 µg/mL). Bacterial extracts showing higher bioactivity (>80% inhibition *in vitro*) were selected for detached-leaf assay against *P. parasitica* on tobacco. In detached-leaf assay, application of 1% ethyl acetate bacterial extract of MOSE L-RD3, MOSEL-RD23, MOSEL-FLS1, MOSEL-FLS3 and MB322 reduced lesion sizes and lesion frequencies caused by *P. parasitica* by 68 to 81%. Over all *P. taiwenensis* MOSEL-RD23 showed positive activities for all the assays. Analysing the potential of bacterial endophytes as biological control agents can potentially lead to the formulation of broad-spectrum biopesticides for sustainable production of crops.

## Introduction

Infectious diseases caused by phytopathogens pose a great threat to food production and stability of ecosystem worldwide [1]. Among the well-recognized plant pathogens, *Phytophthora* spp. are the most destructive Oomycetes, including more than 100 species [2]. *Phytophthora parasitica* is a soil borne pathogen with a wide range of host plants [3]. The severity of *P. parasitica* assumed to the feasibility of zoospores dispersal, infecting plant roots and leaves and rapidly germinating via formation of appressorium at the tip of germ tube [4]. Isolates of *P. parasitica* are capable of infecting a numerous plant species, causing mainly black shank to tobacco as a common worldwide disease [4].

Over the years, biological control of plant pathogens has gained importance because it offers ecological and environmental friendly alternatives for controlling crop diseases [5]. Biocontrol against plant diseases is not only an alternative to chemical pesticides, but it may also provide control of diseases that cannot be managed (or only partly) by other control strategies [6]. The use of beneficial microorganisms (biocontrol agents) is considered as one of the most promising methods for rational and safe agronomical practices. Most of the bacterial strains exploited as biocontrol agents belong to the genera *Agrobacterium, Pseudomonas* and *Bacillus* [7] [8]

Bacteria can also thrive as endophytes in various plants and plant parts and have to be adapted to the specific plant environment, which they colonize and therefore, the metabolic potential of endophytes is likely to differ from their soil dwelling counterparts [9]. Endophytes offer a wide range of benefits to plants by promoting growth via direct and indirect mechanisms [10], direct mechanism such as the production and regulation of levels of phytohormones, including indole-3-acetic acid and ethylene [11, 12], whereas indirect mechanism involve decreased disease severity [13], inducing plant defense mechanisms by the production of toxic compounds (antibiosis) in the host plant; producing substances with antifungal and antibacterial activity [14, 15]. Commercial applications of bacterial endophytes are inoculants in agriculture [16] and a source of secondary metabolites for medical applications [17].

An important mechanism of biological control involves the production of secondary metabolites by these endophytes. Endophytes are less explored for their metabolic potential [9]. Many endophytes are members of genera *Pseudomonas, Burkholderia* and *Bacillus* [18], and these genera are well known for the diverse range of secondary bioactive metabolic products [19]. These secondary metabolites with antagonistic activities have been identified as HCN, phenazines of which major ones are phenazines-1-carboxylic acid and phenazine-1-1carboxymide, 2,4-diacetyl phloroglucinol, pyrrolnitrin, Zwittermycin A and kanosamine. Cyclic compounds are also known for their antagonistic activities [20]. Plants produce several classes of phytohormones including auxins, cytokinins, brassinosteroids, gibberellins, abscisic acid, ethylene, jasmonates and strigolactones playing roles in development and stress responses.

The main objectives of this study were to 1) screen potential bacterial endophytes isolated from *Dodonaea viscosa, Fagonia indica, Caralluma tuberculata* and *Calendula arvensis* for their bioactivity against plant pathogens, 2) extract and assay diffusible and volatile antimicrobial compounds from a select set of endophytes against *Phytophthora parasitica*, and, 3) conduct preliminary fractionation of the bioactive metabolites from a select set of endophytes using High Performance Liquid Chromatography.

## Materials and methods

### Molecular identification and 16S biodiversity analysis

Endophytic bacteria were isolated from four different medicinal plants i.e. *Dodonaea viscosa, Fagonia indica, Caralluma tuberculata* and *Calendula arvensis.* Information about the biological control agents used in this study is summarised in Table 1. Endophytic bacteria were identified using 16S rRNA gene sequencing [21].

**Table 1.** Molecular identification of endophytic bacteria from different plants a) *Dodonaea viscosa* b) *Fagonia indica* c) *Caralluma tuberculata* and d) *Calendula arvensis* using 16S rRNA gene sequencing and anti-*Phytophthora* activity of bacterial extracts at different concentrations against *Phytophthora parasitica*

| Strain ID | Source | Top BLAST search hit (NCBI) | % Nt identity | Disc Diffusion Method | % Growth Inhibition |  |  |
| --- | --- | --- | --- | --- | --- | --- | --- |
|  |  |  |  |  | 400 µg/ml | 40 µg/ml | 4 µg/ml |
| MOSEL-RD3 | *Dodonaea viscosa* | *Streptomyces alboniger* | 100.0 | +++ | 91.96 ± 0.70 | 78.14 ± 1.08 | 4.12 ± 1.07 |
| MOSEL-RD23 | *Dodonaea viscosa* | *Pseudomonas taiwanensis* | 100.0 | +++ | 92.11 ± 1.38 | 47.46 ± 2.99 | 9.96 ± 1.79 |
| MOSEL-RD36 | *Dodonaea viscosa* | *Pseudomonas geniculata* | 100.0 | - | 80.54 ± 0.53 | 5.47 ± 0.88 | 3.74 ± 0.61 |
| MOSEL-FLS1 | *Fagonia indica* | *Enterobacter hormaechei* | 100.0 | +++ | 98.76 ± 0.08 | 98.09 ± 0.11 | 34.49 ± 0.88 |
| MOSEL-FLS3 | *Fagonia indica* | *Bacillus tequilensis* | 100.0 | +++ | 98.75 ± 0.05 | 98.27 ± 0.05 | 40.55 ± 0.72 |
| MOSEL-FIL19 | *Fagonia indica* | *Pantoea* sp. | 95.0 | - | 82.05 ± 0.72 | 50.95 ± 0.86 | 7.2 ± 0.80 |
| MOSEL-MIC5 | *Caralluma tuberculata* | *Bacillus flexus* | 100.0 | +++ | 93.93 ± 0.08 | 52.98 ± 1.18 | 8.36 ± 1.35 |
| MOSEL-MIS12 | *Caralluma tuberculata* | *Pseudarthrobacter phenanthrenivorans* | 100.0 | - | 89.57 ± 0.57 | 76.53 ± 0.57 | 66.15 ± 2.11 |
| MB322 | *Calendula arvensis* | *Delftia lacustris* | 93.60 | +++ | 92.32 ± 1.03 | 86.49 ± 1.03 | 34.38 ± 1.41 |

### *In vitro* antimicrobial activity of bacteria against *Phytophthora parasitica*

*Phytophthora parasitica*, which infects a variety of ornamental, vegetable and citrus [22] was used as target plant pathogen for evaluating the antimicrobial activity of the recovered bacterial endophytes. Dual-culture growth inhibition assays were conducted in petri dishes as follows:A 0.5 cm diameter mycelial plug from 5 days old *P. parasitica* culture was placed in the middle of V8 medium, and incubated for 48 h at 25°C. A 10 µl cell suspension (10^5^ CFU/mL) of potential biocontrol bacteria was inoculated at three spots located 3 cm away from the center of *Phytophthora* plugs. Plates were re-incubated at 25°C for 5 days, and growth inhibition zones of *Phytophthora* were measured.

### Effects of volatiles on the growth of *Phytophthora parasitica*

To investigate the effect of volatiles produced by bacterial endophytes on the growth of the *P. parasitica* bipartite petri dishes were used. Bacterial isolates were inoculated into one compartment of the bipartite petri dish containing TSB media, while *P. parasitica* was placed in the other compartment containing V8 media, and plates were tightly sealed with the parafilm and incubated at 25°C [23].

### Extraction of extracellular secondary metabolites from endophytic bacteria

Selected bacterial strains were inoculated into 10 ml TSB (Oxoid) and incubated for 24 h at 30 °C with shaking until the optical density (OD_600_) reached approximately 0.8 – 1. The culture broth was transferred to 500 mL TSB contained in a 1 L flask and incubated at 30°C in a shaker incubator (220 rpm) for 72 h. At this point, all cultures grew to early stationary phase (OD_600_ = ∼3.0). Bacterial culture was transferred to falcon tubes and cells were pelleted by centrifugation at 8,000 × *g* for 20 min at room temperature. The supernatant was filtered through a 0.20-μm filter and extracted with an equal volume of ethyl acetate as described previously [24] with some modifications. The solvent was then evaporated through vacufuge at 45°C to get the crude extract of bioactive metabolites. The extract was then dissolved in 50 mL methanol.

### *In vitro* antimicrobial activity of bacterial extract

Antimicrobial activity of crude extracts of the selected endophytic bacteria was determined using the disc diffusion method [25]. Sterile filter paper discs were placed approximately 1 cm away from the growing edge of a 24-hr old *P. parasitica* culture on V8 plates. Discs were saturated with each bacterial extract; Methanol was used as a negative control.

### 96-well plate antimicrobial assay of bacterial extract

*In vitro, P. parasitica* growth inhibition assay was performed in 96-well microtiter. Bacterial crude extracts were diluted appropriately 2% to 0.02% in 100 µl sterile water. To each well, 100 µl of freshly prepared *P. parasitica* zoospore suspension (10^5^ zoospores/mL) were added to each well. Each treatment was replicated three times, and each experiment was repeated at least two times. Plates were wrapped with parafilm and incubated at 25°C under continuous light for 18 hrs. Phytophthora growth was quantified using a rapid rasazurine assay (Fai and Grant, 2009, Chromophere 74:1165). At the end of the 18-hr incubation period, 10 µL of a 10% resazurin (Alamar Blue, Cat# DAL1025, Invitrogen) solution was added to each well. Plates were incubated for another 4 hours and the absorbance of the plate was read at 560 and 590 nm wavelength with the Synergy H1 hybrid multimode plate reader (BioTek).

### Bioactivity-driven fractionation of bacterial extracts using HPLC

Fractionation was performed using HPLC according to a method described previously [26]. Chromatographic separation of metabolites was conducted using C18 column using mobile phases A (water with 0.1% formic acid) and B (acetonitrile with 0.1% formic acid) at a flow rate of 0.4 mL/min. HPLC profile consisted of an initial gradient composition (95% A, /5% B) held for 1 min, increased to 20% B over the next 5 min, increased to 80% B over the next 10 min, increased to 100% B over the next 10 min, and held at 100% B for 2 min. For reuse, the initial gradient composition (95% A, /5% B) was restored and allowed to equilibrate for 5 min. The absorption spectrum of fractions was determined in the UV-visible region (230–800 nm) using Synergy H1 Hybrid Multimode spectrophotometer (BioTek). For *in vitro* anti-phytophthora activity of fractions, growth inhibition assay was performed with a 2% final concentration of each fraction in microtiter plates using the rasazurine assay as described above.

### Detached-leaf *Phytophthora parasitica* inhibition assays

The efficacy of bacterial extracts was assessed in a detached leaf assay as described previously [25]. Healthy leaves of tobacco plants from greenhouse-grown were immersed in 15% Chlorox for 2s and immediately rinsed with sterile water three times. Leaves were blot dried lightly and arranged randomly on two layers of moist Whatman filter papers in humid chambers. Each leaf was inoculated with three treatments: 1) Bacterial extract and *P. parasitica* together, 2) methanol & *P. parasitica* (control) and 3) V8 & *P. parasitica* (control).

## Results

### Isolation and Molecular Identification of Bacterial Endophytes

A number of different bacteria were isolated from four different medicinal plants i.e. *Dodonaea viscosa, Fagonia indica, Caralluma tuberculata* and *Calendula arvensis*. On the basis of initial screening assays including cell wall degrading enzymes, siderophore production and antifungal assays, a total of 9 bacteria were selected for further analysis. Based on 16S rRNA gene sequencing, these bacterial isolates were classified into Actinobacteria, Proteobacteria and Firmicutes in nine different genera i.e. *Streptomyces, Pseudomonas, Enterobacter, Bacillus, Pantoea, Pseudarthrobacter* and *Delftia.* Details of nucleotide BLAST searches of these endophytic bacteria are provided in Table 1. Endophytic bacterial isolates with 100% nucleotide identity to known bacterial species were MOSEL-RD3, MOSEL-RD23 and MOSEL-RD36 from *D. viscosa*, MOSEL-FLS1 and MOSEL-FLS3 from *F. indica*, and MOSEL-MIC5 and MOSEL-MIS12 from *C. tuberculata*. Isolates MOSEL-FIL19 from *F. indica* and MB322 from *C. arvensis* showed 95% and 93.6% nucleotide identity to *Pantoea* sp. and *Delftia lacustris*, respectively.

### Screening of endophytic bacteria against *P. parasitica*

Antimicrobial activity of selected bacterial isolates was evaluated against *P. parasitica in vitro*. Visual comparison of assay plates to control plates showed clear growth inhibition by several endophytes as is shown in Figure 1A. Among all the selected isolates, *Pseudomonas taiwanensis* MOSEL-RD23 showed the highest (55%) activity against *P. parasitica* (Fig. 1A and B). *Delftia lacustris* MB322 and *Streptomyces alboniger* MOSEL-RD3 also showed inhibition but to a lesser extent. The remaining isolates showed no or very little activity.

**Figure 1.**
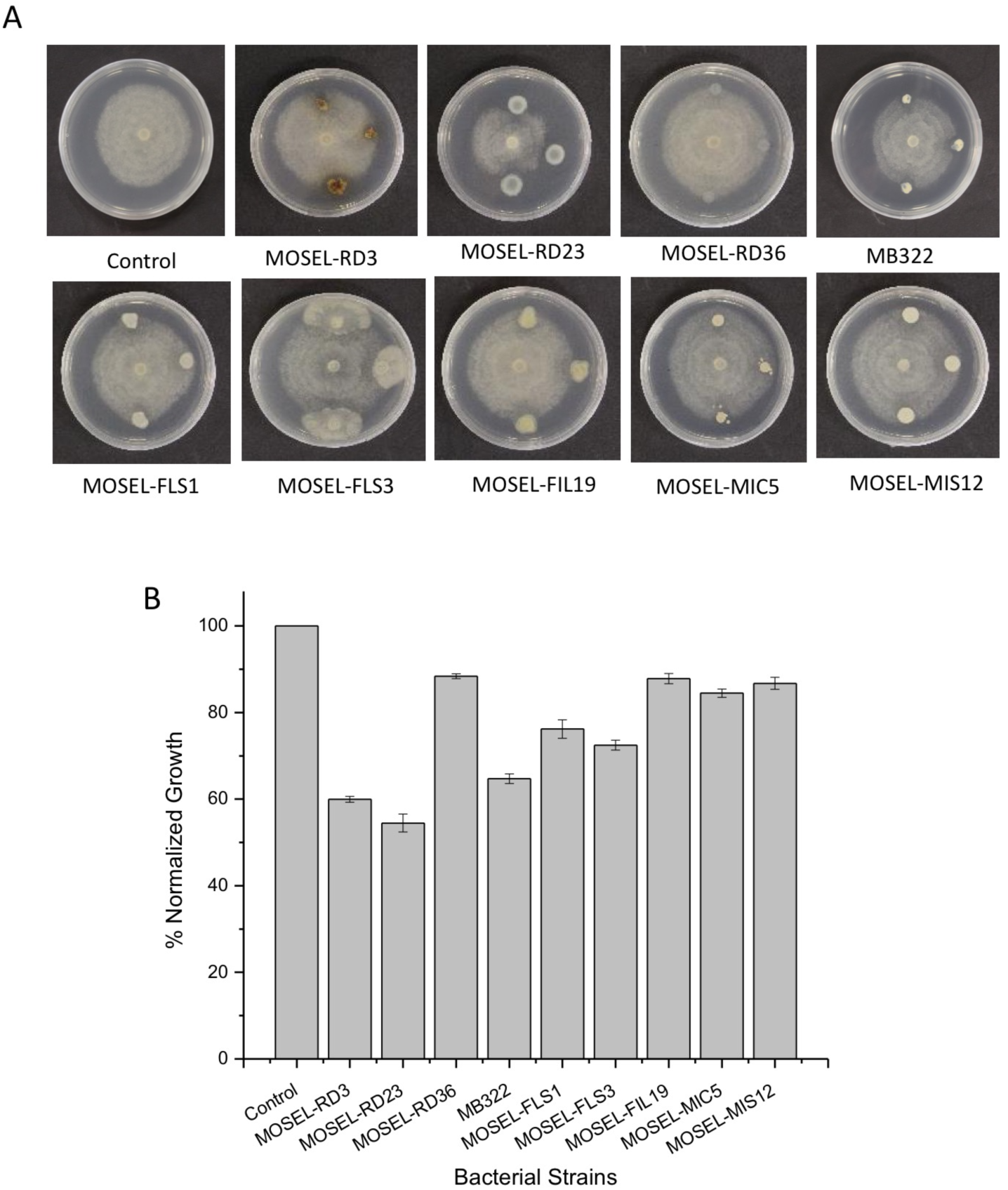
Biological activity of endophytic bacteria against *Phytophthora parasitica.* (A) Representative pictures of plates showing effect of live bacteria on the growth of *P. parasitica.* (B) Bar graphs showing inhibition of *P. parasitica* by endophytic bacteria normalized to growth on control plates. Error bars represent S.E. of means.

### Effect of volatiles on the growth of *P. parasitica*

Visual assessment of initial screening assays revealed that several endophytes slowed growth of *Phytophthora* without showing a clear growth inhibition zone. This observation suggested that these endophytes might be emitting volatile compounds that are inhibitory to *Phytophtora*. To test this hypothesis, we conducted bipartite split-plate growth inhibition assays, in which *Phytophthora* was physically separated from potential antimicrobial endophytes but was otherwise exposed to gaseous antimicrobial compounds emitted by endophytes. These assays showed that the growth of the *P. parsitica* was significantly inhibited in the presence of several bacterial strains. A majority of bacteria showed growth inhibition at varying levels ranging from about 50 to 80% reduction as compared to control (Fig. 2A & B). *Pseudomonas taiwnensis* MOSEL-RD23, *Bacillus flexus* MOSEL-MIC5 and *Streptomyces alboniger* MOSEL-RD3 were the most effective, displaying 50 - 60% growth inhibition. The remaining isolates, *Pseudarthrobacter phenanthrenivorans* MOSEL-MIS12, *Enterobacter hormaechei* MOSEL-FLS1, *Bacillus tequilensis* MOSEL-FLS3, *Pantoea* sp. MOSEL-FIL19 *Delftia lacustris* MB322 and *Pseudomonas geniculata* MOSEL-RD36 showed some activity resulting in less than 50% reduction in mycelial growth as compared to the control.

**Figure 2.**
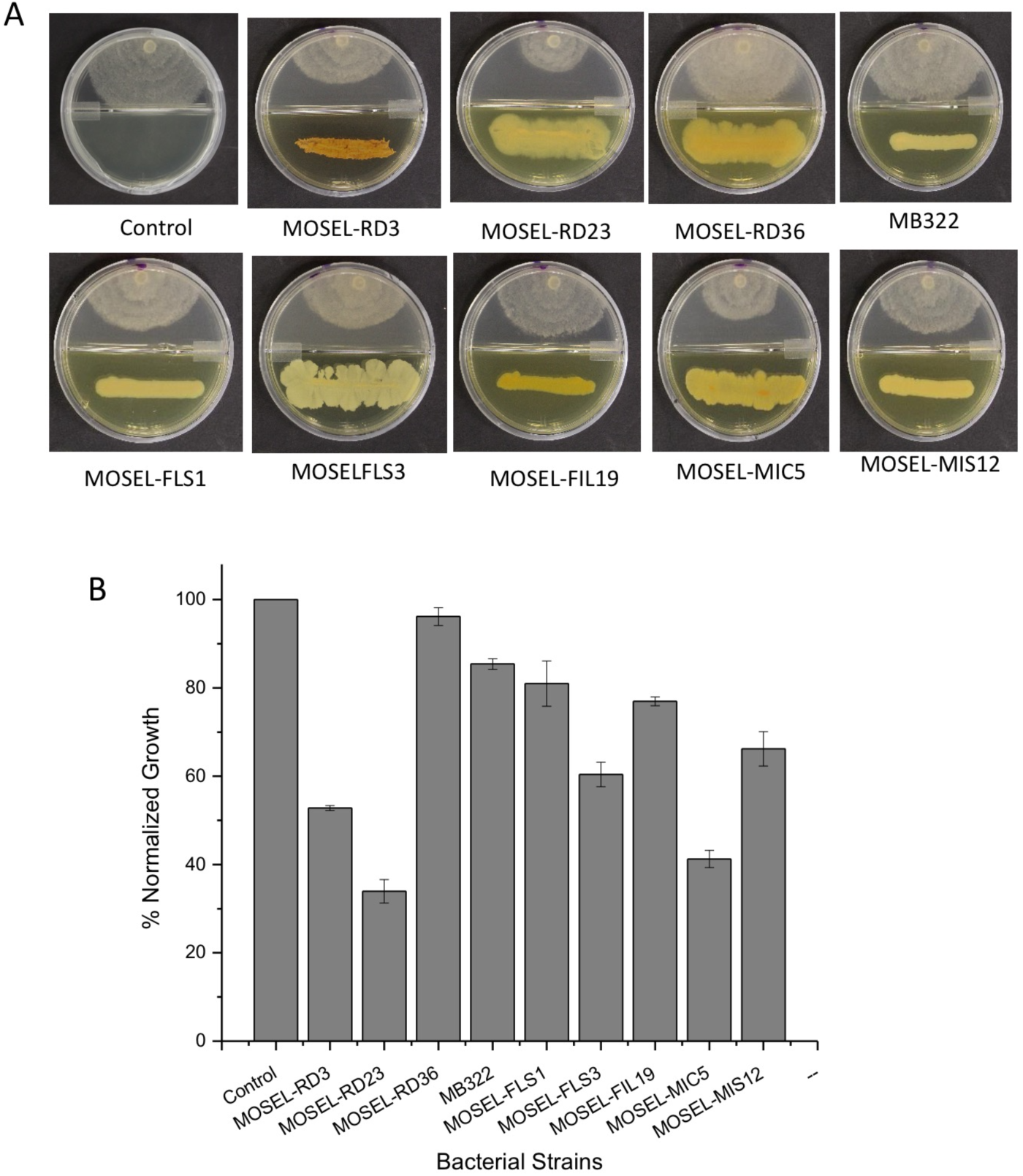
Effect of volatiles produced by endophytic bacteria on the growth of *Phytophthora parasitica.* (A) Representative pictures showing effect of volatiles produced by endophytic bacteria on the growth of *P. parasitica*. (B) Bar graph showing % normalized growth of *P. parasitica* as a result of production of volatiles by endophytic bacteria.

### Assaying extracellular secondary metabolites for inhibiting growth of *P. parasitica*

Bioactivity of ethyl acetate extract from endophytic bacteria culture supernatant was assessed using the disc diffusion method against *P. parasitica*. Bacterial extracts of *S. alboniger* MOSEL-RD3, *P. taiwanensis* MOSEL-RD23, *Delftia lacustris* MB322, *E. hormaechei* MOSEL-FLS1 and *B. tequilensis* MOSEL-FLS3 showed very strong activity against *P. parasitica* (Fig. 3A). The remaining isolates showed no or very little activity. These bacteria displayed more or less similar growth inhibition ranging from 40 to 45% (Figure 3B). The rest of the isolates were less effective displaying 20 to 30% growth inhibition. Microscopic examination of the growth inhibition zones showed reduced and scattered mycelial growth compared to growth on control plates (Fig. 3B). Zone of inhibition was measured in terms of growth inhibition normalized to growth on control plates (Fig. 3C).

**Figure 3.**
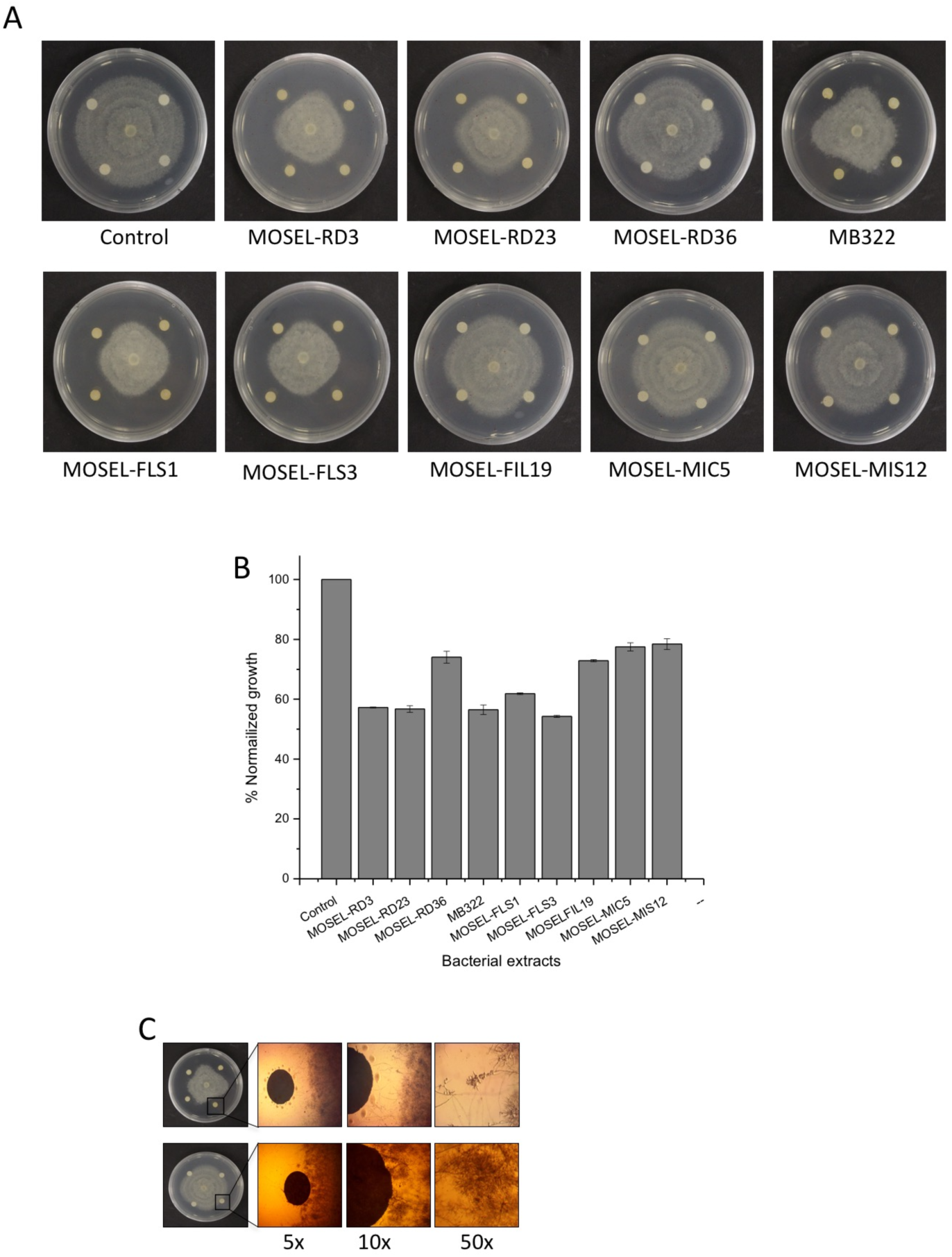
Bioactivity of ethyl acetate extract from endophytic bacteria culture supernatant showing inhibition against *Phytophthora parasitica.* (A) Bacterial extracts (MOSEL-w2, MOSEL-w7, MOSEL-w13, MB322, MOSEL-RD3, MOSEL-RD23, MOSEL-RD36, MOSEL-FLS1, MOSEL-FLS3 and MOSEL-FIL19) showing strong activity against *P. parasitica*. (B) Representative micrographs of bacterial extract showing inhibition of *P. parasitica* as compared to control at different magnification (7.5, 10 and 15) under compound microscope. (C) Bar graph showing growth inhibition of *P. parasitica* by bacterial extracts normalized to growth on plates

Bioactivity of the ethyl acetate extract of the culture supernatants from endophytic bacteria against *P. parasitica* was assessed at different concentrations using resazurin-based antimicrobial assays in microtiter plates. Ethyl acetate extracts dissolved in methanol were tested using three 10-fold serial dilutions at 2 % (400 µg/ml), 0.2% (40 µg/ml) and 0.02 % (4 µg/ml). At the 2% concentration levels, extracts from all bacteria showed strong but mostly similar growth inhibition (>80%) against *P. parasitica*. Similarly, at the 0.2% level, ethyl acetate extracts from all bacteria also showed growth inhibition. However, they differed significantly in the levels of inhibition with MOSEL-RD3, MB322, MOSEL-FLS1 and MOSEL-FLS3 showing strong (> 80%) growth inhibition, whereas MOSEL-RD23, MOSEL-FIL19, MOSEL-MIC5 and MOSEL-MIS12 showing intermediate (approx. 50%) inhibition. Inhibition assays with the lowest concentration (0.02%) showed that the extracts of MB322 still displayed strong inhibition against *P. parasitica* (Table 1; Figure 4A). Micrographs of hyphae of *P. Parasitica* treated with the selected bacterial extracts showed abnormal growth compared to control (Figure 4B). Representative pictures showing convoluted, swollen nodes and abnormal growth of hyphae of *P. parasitica* in response to bacterial extracts are shown in Figure 4C.

**Figure 4.**
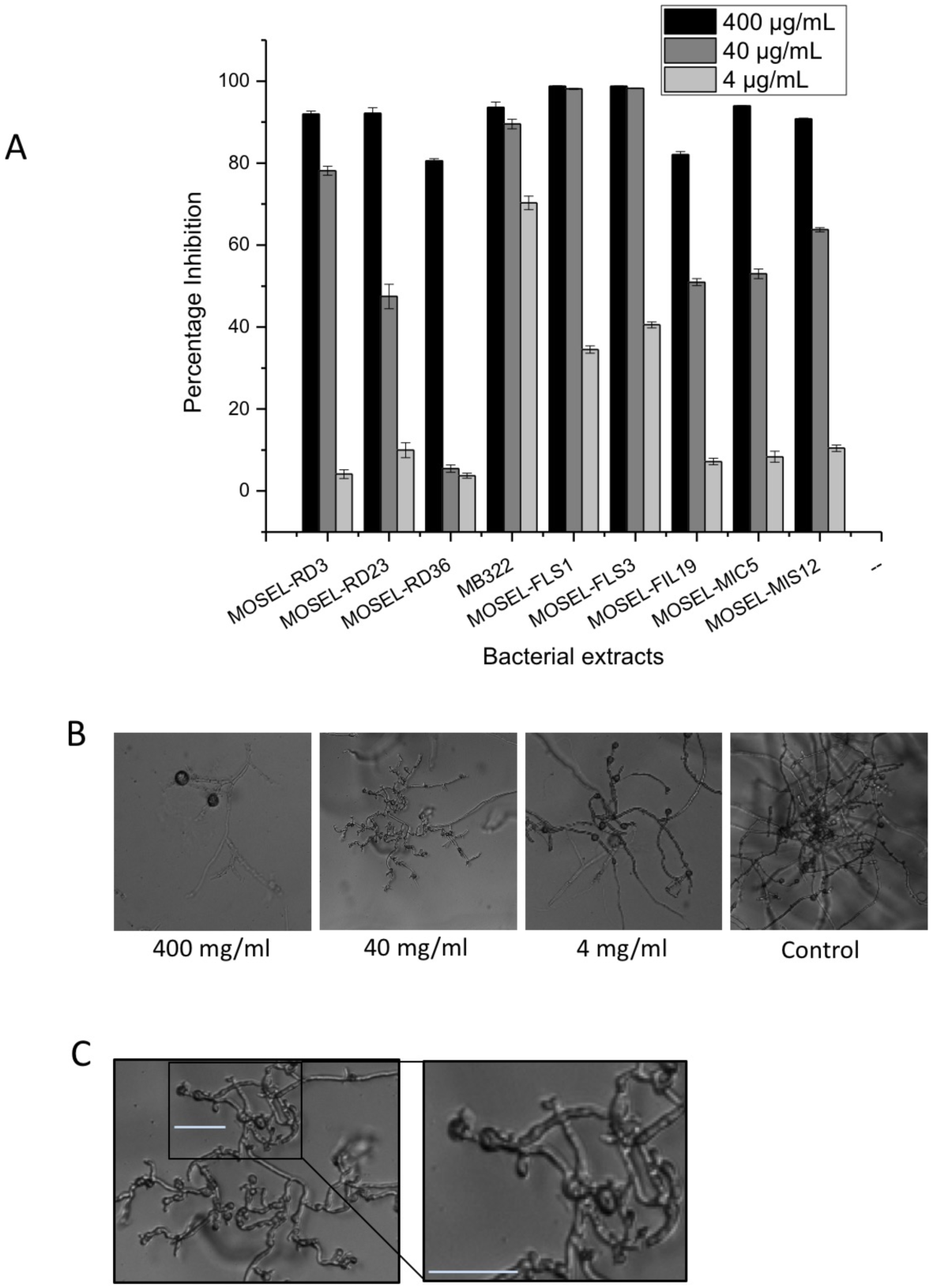
Bioactivity of the ethyl acetate extract from endophytic bacteria culture supernatant at different concentrations against *Phytophthora parasitica.* (A) Bacterial extracts showing %inhibition of *P. parasitica* at different concentrations a) 400 µg/ml b) 40 µg/ml and c) 4 µg/ml. (B) Micrographs showing abnormal growth of hyphae of *P. Parasitica* in response to the different concentrations of bacterial extracts as compared to control. (C) Micrograph showing convoluted, swollen nodes and abnormal growth of hyphae of *P. parasitica* in response to bacterial extract

### Detached-leaf *P. parasitica* inhibition assays

The *in vivo* anti-*Phytophthora* activity of bacterial extracts were determined using a tobacco detach leaf assay. In this assay, bacterial extracts of the selected bacterial strains were tested for preventing expansion of lesions caused by *P. parasitica* on tobacco leaves. As is shown in Figure 5A and 5B, all bacterial extracts inhibited expansion of lesions produced by *P. parasitica* on tobacco leaves as compared to negative controls, which consisted of methanol only plus *P. parasitica* or V8 plus *P. parasitica.* Tryphan blue staining of inoculated leaves showed that in the control treatments, Phytophthora had grown extensively beyond the inoculation spots. In contrast, very little growth was observed outside the spots treated with ethyl acetate bacterial extracts (Figure 5A). Quantification of lesion sizes showed an average of 50% reduction by bacterial extracts compared to controls (Figure 5B). Overall, these results indicate that ethyl acetate extracts of these bacteria contain antimicrobial properties, which are active *in vivo*.

**Figure 5.**
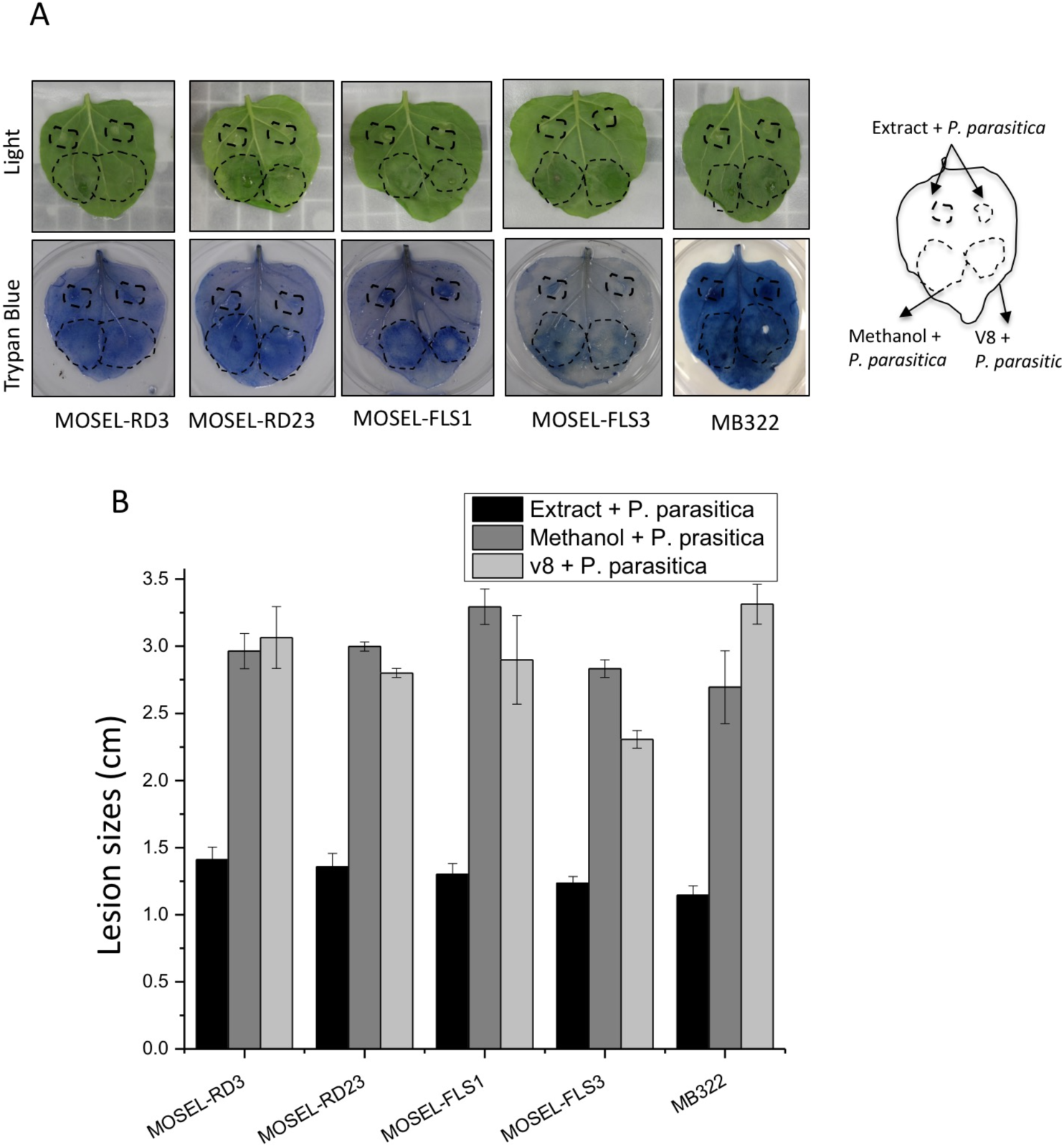
In vivo detached lead Bioactivity of 1% bacterial extracts against *Phytophthora parasitica.* (A) Detached leaf assay showing activity of bacterial extract of MOSEL-RD3, MOSEL-RD23, MOSEL-FLS1, MOSEL-FLS3 and MB322 on the growth of *P. parasitica.* (B) Bar graphs showing average lesion diameter caused by *P. parasitica* on tobacco leaves

### Bioactivity-driven fractionation and UV-Vis scanning

Preparative HPLC was used to fractionate culture filtrates for finding fractions with bioactivity. All fractions of each extract were assayed for their bioactivity against *P. parasitica* in vitro. These analyses showed variable bioactivity in multiple fractions (Figure 6). Although, none of the fractions of any of the bacteria stood out as the sole source of anti-*Phytophthora* activity, in general, MB322 and MOSEL-FLS1 exhibited a trend of increasingly stronger activity in fractions with longer retention times. In contrast, MOSEL-RD3 displayed stronger activity in fractions with early retention time points, starting at 7 min, which then displayed a declining trend towards the end of the run. Mosel-RD23 did not reveal a defined trend of anti- *Phytophthora* activity with almost all fractions showing strong activity except a few fractions in the middle of the run, which showed very little activity. MOSEL-FLS3 displayed very different but a well-defined trend with fractions with retention times between 15 to 27 minutes displaying a normal distribution. These differential results show that the tested isolates most likely have antimicrobial compounds with different biochemical properties. Most fractions that displayed strong activity also caused different abnormal growth patterns such as convoluted hyphae, swollen nodes, and constricted and excessive branching, suggesting that inhibition of *P. parasitica* is associated with effect on different cellular and physiological processes.

**Figure 6.**
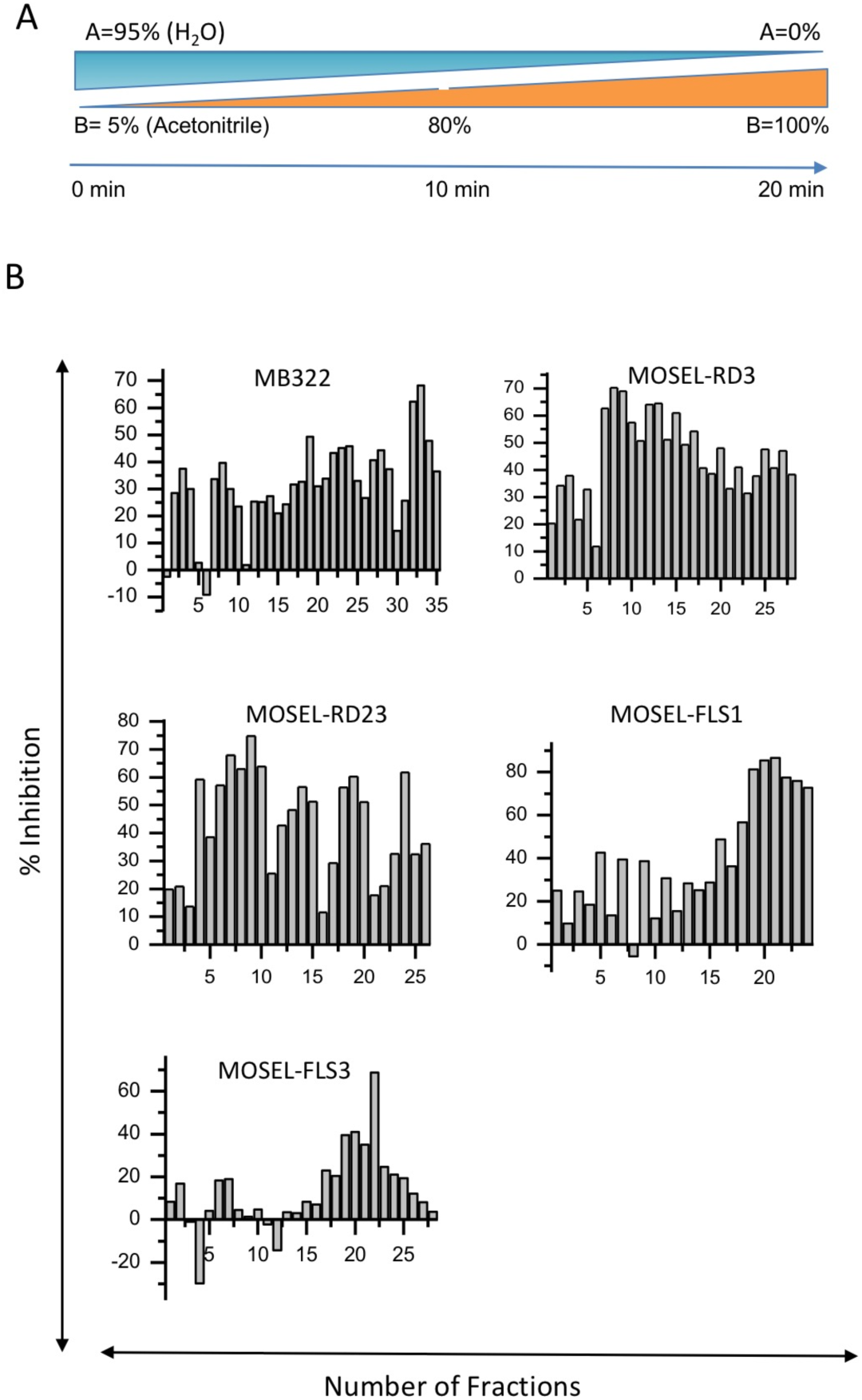
Bioactivity of fractions of the ethyl acetate extract from selected endophytic bacteria culture supernatant obtained through high-pressure liquid chromatography (HPLC) against *P. parasitica*

In order to determine the absorbance properties of bacterial extracts, fractions of two bacterial isolates, MOSEL-FLS1 and MOSEL-FLS3, which displayed different HPLC-determined antimicrobial patterns, were subjected to UV-Vis spectral scanning at 230 – 1000 nm. These analyses revealed that different fractions of both bacterial isolates contained compounds with strong absorbance in the 230 to 290 nm range (Figure S3). However, consistent UV-Vis patterns with any significant correlation with the HPLC-based inhibition patterns were not observed. These results further suggested that antimicrobial compounds in different fractions of different bacteria have different physicochemical properties.

## Discussion

Medicinal plants usually grow vigorously in the wild without any significant irrigation, and fertilizer and pesticides inputs. Association with beneficial microbes is considered as one of the major contributors to the enhanced growth of these medicinal plants [27]. However, microbiomes of medicinal plants are less explored. Medicinal plants used in this study are well-known for their therapeutic potential. *Dodonaea viscosa* is an evergreen, woody and wild shrub growing throughout the tropics. It is found in sub-tropical regions and is known for its medicinal value [28]. *Fagonia indica* is a small spiny shrub with reputed medicinal value and is distributed in the warm and dried areas of the middle east and the Indian subcontinent [29, 30]. *Calendula arvensis*, commonly known as Field Marigold, is an annual herb, which is naturally distributed in Southern Europe, Iran, Afghanistan and Pakistan and is known for various ethnic medicinal properties [31, 32]. *Caralluma tuberculata* is a herbaceous, succulent and leafless plant, which is naturally found in the dry areas of Africa, South Mediterranean and tropical Asia and it is also known for several medicinal properties [33, 34]. Similar to many medicinal plants, the microbiome of these plants is also not known. In this report we investigated several endophytic bacteria isolated from these plants for anti-Phytophthora activities.

Initial characterization of bacteria isolated from these four plants showed that several of them produce cell wall degrading enzymes and siderophores, which have previously been associated with plant growth promotion and antimicrobial activities (ref). Endophytic bacteria isolated from these plants were explored for their potential as biocontrol agents. *Streptomyces alboniger, Pseudomonas taiwanensis* and *Pseudomonas geniculata* from *Dodonaea viscosa*, which were previously reported as potential plant growth promoting bacteria [21]. *Enterobacter hormaechei* and *Bacillus subtilis* has been suggested to play a role in the therapeutic properties of *Fagonia indica* [35]. *Bacillus flexus* and *Pseudarthrobacter phenanthrenivorans, Delftia and Baciluus tequilis, which all displayed very potencies*, are not widely studied for their biological control potential especially against *Phytophthora* disease. Genome of type strain of *Pseudarthrobacter phenanthrenivorans* Sphe3, which was isolated from a creosote-polluted site in Greece has been sequenced [36]. Similarly, genome of another strain, isolated from arsenic-contaminated soil in China has also been sequenced [37]. Sequence annotation of these strains revealed genes involved in pollution detoxification and tolerance particularly against arsenate and several metals[36, 37]. They are, however, not investigated extensively for their antimicrobial activity. Nevertheless, genes associated with the synthesis and regulation of several polypeptides have been reported, some of which might be involved in the anti-Phytophthora activity of the isolate reported here [36, 37]. Whether or not the strain reported in this study contain genes for and metabolic pathways for production of compounds and enzymes that might be involved in inhibiting the growth of Phytophthora remains to be investigated. We have planned on sequencing the genome of the isolate of *Pseudarthrobacter phenanthrenivorans* reported in this study, which will provided clues about its mode of action. Since *Pseudarthrobacter phenanthrenivorans* isolates have not been studied extensively or commercialized as biological control agents, genome information and anti-*Phytophthora* studies reported here will be useful in the development of this strain into a commercial biopesticide. A new biological control agent with anti-Phytophthora activity will provide disease management tool for organic production of crops by suppressing soil-borne Phytophthora, which are important plant pathogens causing significant economic and yield losses in a wide range of crops [38].

In this report, we screened the above-mentioned bacteria for their potential application in the biological control of Phytophthora diseases. In in vitro growth inhibition assays, *Pseudomonas taiwanensis* MOSEL-RD23 exhibited the highest antifungal activity against *P. parasitica*. *Azotobacter* sp., *Pseudomonas* sp., *Streptomyces* sp. and *Bacillus* sp. have previously been reported for antifungal activity [39]. These bacterial endophytes help in plant growth promotion and in control of pathogens. Positive effects displayed by bacterial endophytes are mediated by several metabolites acting together [9]. Similar to previous studies, bacterial extracts of biocontrol agents reported in this study also induced abnormal growth in the hyphae of *P. parasitica*. These anomalies are most likely due to compounds that negatively affect hyphal growth [40].

Diffusible and volatile compounds both showed activity against Phytophthora suggesting that the biological control agents reported in this study employ various modes of actions. Bioactivity-driven fractionation of ethyl acetate extracts of endophytic bacteria using HPLC analyses showed anti-Phytophthora activity is not restricted to a specific fraction. Strikingly, almost all fractions showed activity against Phytophthora suggesting that antimicrobial compounds responsible for controlling Phytophthora contain different physical and chemical properties. This is also consistent with previous reports where different compounds such as lytic peptides, antibiotics and other secondary metabolites contribute to overall biocontrol activity of these biocontrol agents (reviewed in [41]). These extracts might have soluble and volatile compounds, which, in future studies will be further characterized using LC-MS/MS. These future studies could potentially result in the identification of compounds with different modes of actions, which could be utilized in formulating synthetic blends of biocontrol agents for optimal plant growth promotion and disease suppression.

## Conclusions

This is the first report about the biological activities of secondary metabolites as biocontrol agents.

## Acknowledgements

This work was supported by funding from Higher Education Commission of Pakistan to I.I. and by the National Institute of Food and Agriculture – United States Department of Agriculture (Accession number 1017239 and FL-APO-005155) to G.A.

